# KMT5C-H4K20me3 drives changes in epigenetic landscape independent of H3K9me3

**DOI:** 10.64898/2026.05.11.724304

**Authors:** Jihye Son, Ching-Hua Shih, Christina Davidson, Sagar M. Utturkar, Nathaniel W. Mabe, Alexandra Glaws, Paula M. Vertino, Andrea L. Kasinski

## Abstract

Histone H4 lysine 20 trimethylation (H4K20me3) is a histone modification that is critical in maintaining genome integrity. Dysregulation of H4K20me3 and KMT5C, the major methyltransferase for H4K20me3, occurs commonly in multiple types of cancer but the mechanisms surrounding how they contribute to shaping the epigenomic landscape remains unclear. Here, we show that KMT5C is involved in non-canonical deposition of H4K20me3, independent of H3K9me3, which was previously recognized as a prerequisite for H4K20me3. This novel subtype of H4K20me3 lacks canonical repressive epigenetic signatures and instead overlaps with multiple activating marks. These activating modifications likely contribute to the dynamic changes in transcript levels upon loss of H4K20me3. The mechanism involved in recruiting KMT5C to these loci is independent of HP1, the factor reported to be involved in recruitment of KMT5C to heterochromatin marked with H3K9me3. Instead, biochemical analyses revealed ZNF280C to be a novel interacting partner of KMT5C, with ZNF280C localizing specifically at H3K9me3-/H4K20me3+ sites. Together, these results suggest a novel, non-canonical function of KMT5C-H4K20me3 that protects vulnerable regions of the genome from uncontrolled expression.

## INTRODUCTION

Post-translation modifications (PTM) of histones dynamically regulate chromatin-based processes that can alter accessibility to transcription machinery and expression of genes under their influence [1]. Dysregulation of enzymes that are responsible for histone PTMs is expected to occur in 10-20% of all cancers [2], and their role in tumorigenesis has been recognized as a novel hallmark of cancer [3,4]. In support of this, we recently reported that loss of KMT5C, a methyltransferase for histone H4 lysine 20 trimethylation (H4K20me3), promotes resistance to EGFR inhibitors in EGFR-mutant non-small cell lung cancer [5]. Others have also independently demonstrated that KMT5C-H4K20me3 is dysregulated in multiple types of cancer, including lung cancer [6–11]. Nevertheless, a thorough investigation of how KMT5C-H4K20me3 contributes to the regulation of gene expression has not been conducted. Elucidation of such mechanisms are integral to understand how KMT5C-H4K20me3 is involved in tumor initiation, progression, and drug resistance.

KMT5C is reported to be recruited to chromatin in a H3K9me3-dependent manner through the interaction between KMT5C and heterochromatin protein 1 (HP1), which itself associates with H3K9me3. Following HP1 localization at heterochromatin marked with H3K9me3 through its chromodomain, HP1 interacts with KMT5C through the C-terminal domain (CTD) of KMT5C [12– 14]. While this interaction has been reported to be reproducible in constitutive heterochromatic regions where the majority of H3K9me3 resides, several groups, including our own, have reported that KMT5C-H4K20me3 is also present in regions outside of heterochromatin [5,15]. Evidence from *in vitro* models further suggests that KMT5C retains the ability to affect expression of its targets independent of its interaction with HP1 [16]. This raises the possibility that additional mechanisms, other than HP1-mediated recruitment of KMT5C to the chromatin, could be influencing KMT5C localization.

Here, we aimed to comprehensively define the deposition of KMT5C-mediated H4K20me3 across gene-rich regions, including regions that are either constitutively repressed and marked with repressive histone PTMs or poised for expression and associated with activating histone PTMs, and to determine how gene expression changes upon perturbation of H4K20me3. Our work uncovered a previously unreported, non-canonical function of KMT5C, whereby KMT5C can deposit H4K20me3 independently of H3K9me3. Genes lacking H3K9me3 were particularly sensitive to the loss of H4K20me3. To investigate how KMT5C is recruited to these regions, we employed a proximity-labeling BioID approach, which revealed a previously unidentified interaction between KMT5C and ZNF280C at H3K9me3-/H4K20me3+ loci that may contribute to transcriptional regulation of genes with this signature. Collectively, these findings establish a novel, non-canonical role for KMT5C in shaping the epigenomic landscape independently of H3K9me3, thereby regulating a subset of genes poised for expression.

## RESULTS

### Identification of a non-canonical H4K20me3 subclass that is devoid of H3K9me3

In addition to its correlation with repressive histone modifications, several studies have demonstrated that H4K20me3 is deposited outside of heterochromatic regions [5,17,18]. Consistent with this, H3K9me3, previously reported as an upstream requirement for H4K20me3, does not perfectly overlap with H4K20me3 in NCI-60 cell lines, suggesting that other subpopulations of H4K20me3 chromatin exist [15]. To directly assess the extent of co-occupancy, we profiled genome-wide distribution of H4K20me3 and H3K9me3 using <u>c</u>leavage <u>u</u>nder target & <u>r</u>elease <u>u</u>sing <u>n</u>ucleases (CUT&RUN). HCC827 and PC9 cells were used for this work based on our earlier study that determined that H4K20me3 plays a key role in driving resistance to EGFR inhibitors in these cells [5]. As reported previously [12], a subset of H4K20me3 peaks overlapped with H3K9me3. However, 38% of H4K20me3 peaks lacked H3K9me3 enrichment (Figure 1A-C and Figure S1A). Annotation of these H3K9me3-/H4K20me3+ peaks to the nearest transcription start sites revealed enrichment for genes involved in DNA replication, RNA processing, and cell cycle transitions (Figure S1B). The majority of these genes were shared between HCC827 and PC9 cells, suggesting that the H3K9me3-/H4K20me3+ signature represents a conserved epigenetic program (Figure 1D). Validation of select loci by CUT&RUN followed by quantitative PCR (CUT&RUN-qPCR) confirmed that H4K20me3 occurs independently of H3K9me3 (Figure 1E-F). This conclusion was further supported by analysis of published <u>c</u>leavage <u>u</u>nder targets & <u>tag</u>mentation (CUT&Tag) data in A549 (lung cancer) and SK-Mel-127 (melanoma) cell lines [19], which also revealed a subset of H4K20me3 peaks lacking H3K9me3 co-enrichment (Figure S1C-D)[19]. Together, these findings demonstrate that H4K20me3 may be deposited through a non-canonical mechanism.

**Figure 1.**
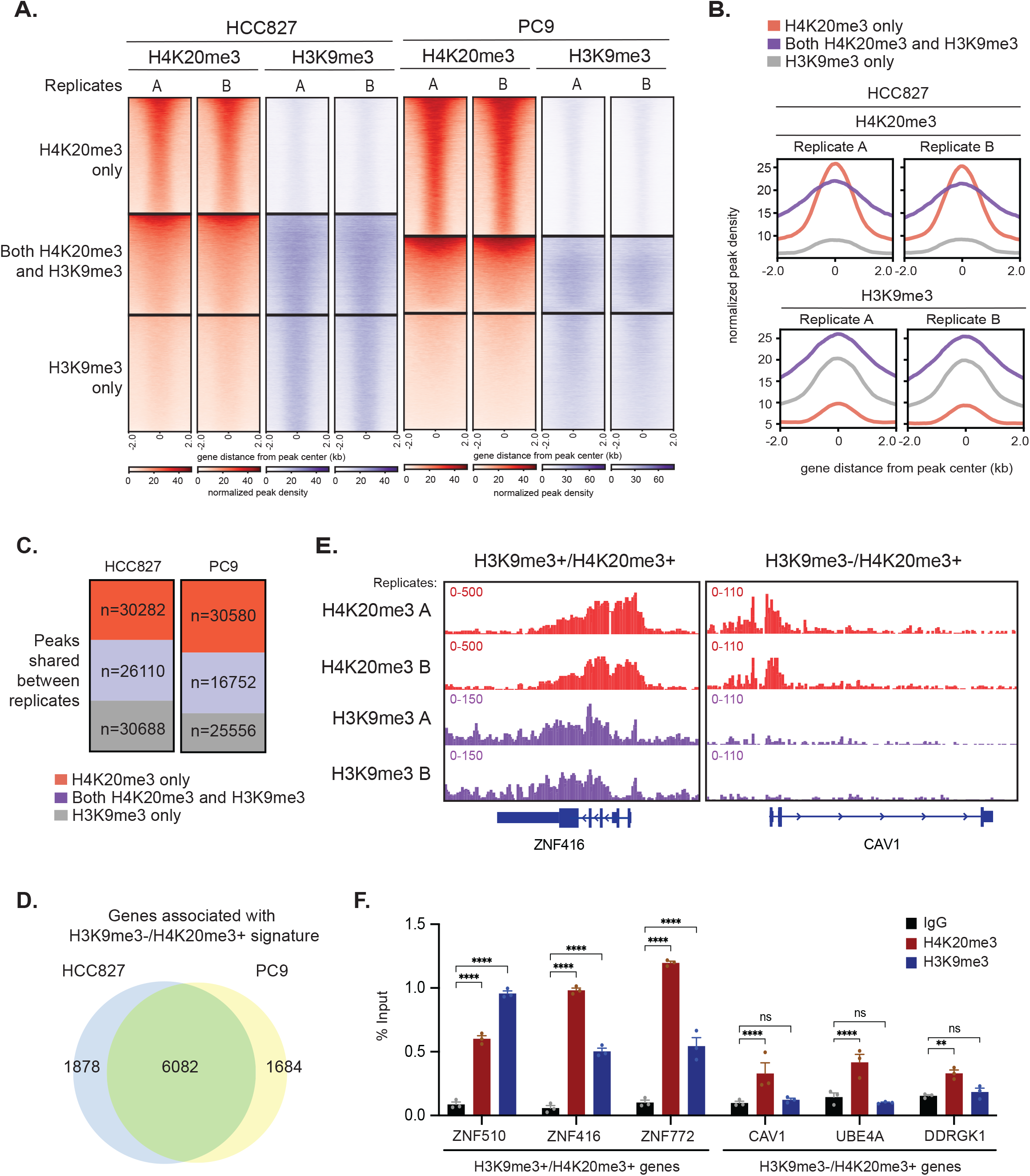
Non-canonical enrichment of H4K20me3 independent of H3K9me3. **A**. Heatmaps of H4K20me3 and H3K9me3 peaks in HCC827 and PC9 cells, clustered by overlap between the two modifications. n=2 biological replicates. **B**. Normalized peak density plots of H4K20me3 and H3K9me3 peaks in HCC827 cells, clustered by the overlap between the two modifications. **C**. Counts of H4K20me3-only, H3K9me3-only, and overlapping H4K20me3/H3K9me3 peaks shared between biological replicates in HCC827 and PC9 cells. **D**. Number of genes exclusively associated with the H3K9me3-/H4K20me3+ signature in HCC827 and PC9 cells. **E**. IGV browser tracks showing representative genes with H3K9me3-/H4K20me3+ peaks (ZNF416) and H3K9me3+/H4K20me3+ peaks (CAV1) in HCC827 cells. **F**. CUT&RUN-qPCR validation of selected loci in HCC827 cells. Data shown are representative technical replicates from one of three biological replicates. **P<0.01, ***P<0.001, ****P<0.0001

### The H3K9me3-/H4K20me+ signature lacks canonical repressive epigenetic marks

To determine whether H3K9me3+/H4K20me3+ and H3K9me3-/H4K20me3+ regions have distinct roles, we examined their genomic location. Whereas H3K9me3+/H4K20me3+ peaks were broadly distributed across gene bodies, the H3K9me3-/H4K20me3+ peaks were concentrated at transcription start sites, suggesting discrete roles for these two subpopulations (Figure 2A and Figure S2A). We next performed genome-wide CUT&RUN profiling of H3K27me3, H3K27ac, H3K4me3, and H4K20me1, and assessed the distribution within each signature. As expected for repressive chromatin, H3K9me3+/H4K20me3+ peaks were generally depleted of activating marks, including H3K27ac and H3K4me3. In contrast, the H3K9me3-/H4K20me3+ subclass displayed co-enrichment of activating modifications and reduced loci marked with H3K27me3, consistent with a more permissive chromatin state. We further confirmed that H4K20me3 co-localizes with KMT5C, the primary H4K20me3 methyltransferase (Figure 2B-C and Figure S2B-C).

**Figure 2.**
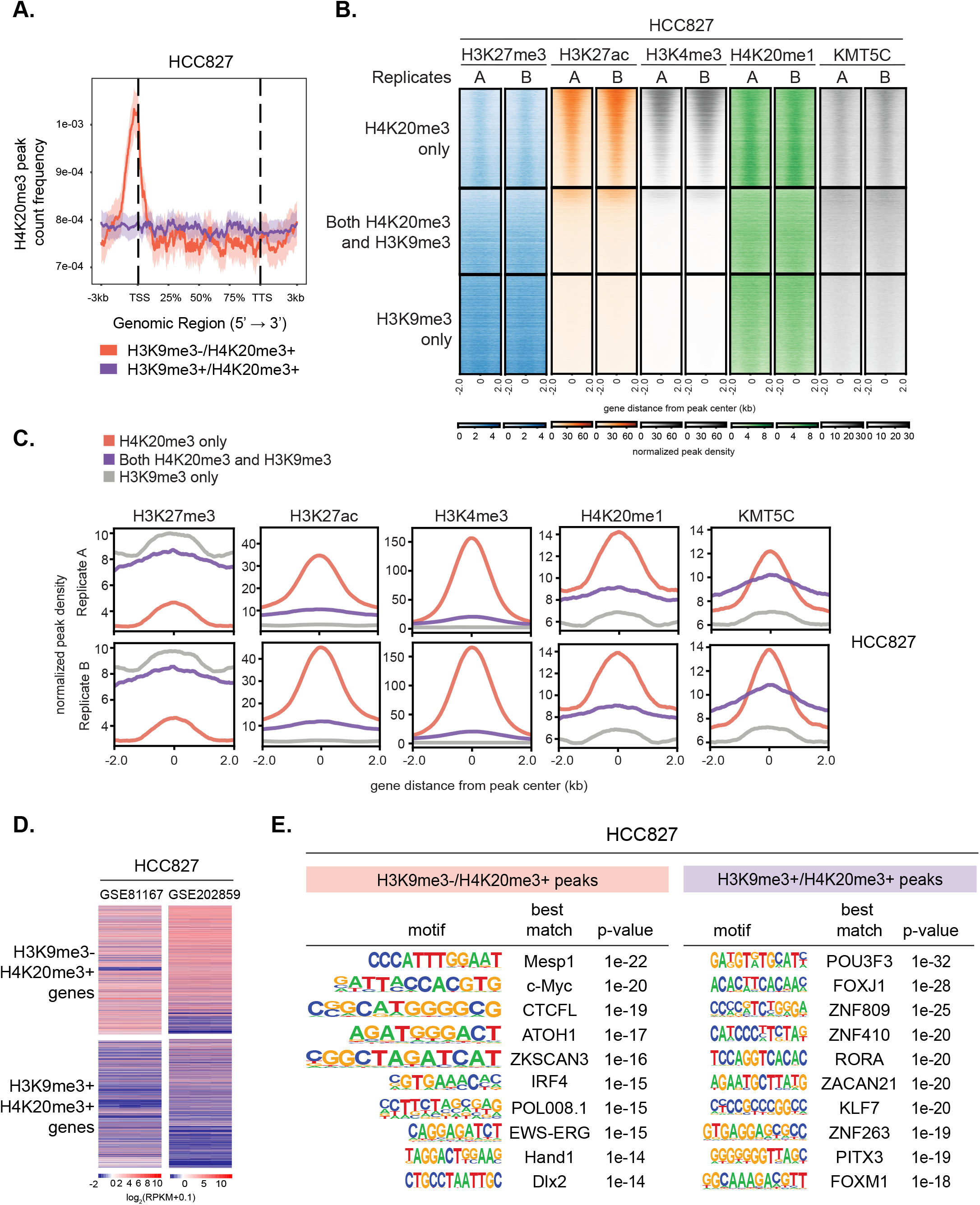
H3K9me3-H4K20me3 peaks lack canonical repressive epigenetic signature. **A**. Distribution of H3K9me3-/H4K20me3+ and H3K9me3+/H4K20me3+ across the gene bodies in HCC827 cells. TSS, transcriptional start site; TTS, transcriptional termination site. **B**. Heatmaps of H3K27me3, H3K27ac, H3K4me3, H4K20me1, and KMT5C peaks in HCC827 cells, clustered by H4K20me3-only, H3K9me3-only, or overlapping H4K20me3/H3K9me3 peaks. **C**. Normalized peak density plots of H3K27me3, H3K27ac, H3K4me3, H4K20me1, and KMT5C in HCC827 cells, clustered as in (B). **D**. Expression of genes associated with H3K9me3-/H4K20me3+ or H3K9me3+/H4K20me3+ peaks in HCC827 cells. RNA-seq data were obtained from GSE81167 and GSE202859. **E**. HOMER motif analysis of H3K9me3-/H4K20me3+ or H3K9me3+/H4K20me3+ peaks in HCC827 cells.

To assess the transcriptional impact of these signatures, we analyzed RNA-seq datasets from HCC827 and PC9 cells (GSE81167, GSE202859, and GSE129221) [20–22]. Genes associated with the H3K9me3-/H4K20me3+ signature were expressed at higher levels than those associated with H3K9me3+/H4K20me3+ peaks (Figure 2D and Figure S2D). In the context of the full transcriptome, loci with H3K9me3-/H4K20me3+ were among the most highly expressed, highlighting that H3K9me3-/H4K20me3+ signature contributes to a more permissive chromatin (Figure S3A-B). Motif enrichment analysis further revealed distinct sequence preferences between the two signatures. Notably, MYC-related motifs were specifically enriched within the H3K9me3-/H4K20me3+ subpopulation in both HCC827 and PC9 cells (Figure 2E and Figure S2E). Together, these findings indicate that H3K9me3–/H4K20me3+ peaks represent a unique chromatin signature characterized by activating histone marks, increased transcriptional activity, and unique sequence motifs, highlighting a potential distinct role for this non-canonical H4K20me3 deposition in chromatin regulation.

### H4K20me3 loss leads to dynamic changes in expression of H3K9me3-/H4K20me3+ genes

To investigate how H4K20me3 influences transcriptional programs, we performed RNA-seq after treatment with A-196, a small-molecule inhibitor of KMT5B and KMT5C, which reduces global H4K20me3 levels (Figure 3A) [23]. Differentially expressed genes following A-196 treatment were associated with various pathways, including those related to epithelial-to-mesenchymal transition (EMT), consistent with prior reports (Figures S4A-B) [7,24]. Notably, a greater fraction of genes associated with H3K9me3-/H4K20me3+ peaks were altered compared to those linked to H3K9me3+/H4K20me3+ peaks (Figure 3B-D and Figures S4C-D). To further assess functional impact, we integrated RNA-seq data with H4K20me3 CUT&RUN profiles from Figure 1 and used BETA analysis to predict the functions of H3K9me3-/H4K20me3+ and H3K9me3+/H4K20me3+ peaks [25]. Even though genes associated with the H3K9me3-/H4K20me3+ subclass were among the most highly expressed (Figure S3), BETA predicted a repressive role for H3K9me3-/H4K20me3+, whereas H3K9me3+/H4K20me3+ peaks were not significantly associated with either repression or activation (Figure 3E). Together, these findings suggest that H4K20me3 at H3K9me3-sites functions as a safeguard to restrain excessive gene expression, such that loss of H4K20me3 at these regions leads to dynamic transcriptional changes. By contrast, H3K9me3+/H4K20me3+ sites appear to harbor multiple safeguards, buffering gene expression against change even when H4K20me3 is reduced. To confirm the role of KMT5C in this process, we performed KMT5C knockdown in HCC827 cells and examined transcripts of candidate genes identified from RNA-seq (Figure S4E). KMT5C knockdown reduced H4K20me3 levels, albeit less effectively than A-196 treatment (Figure 3F, compare A-196 treatment in the presence of siNC to DMSO treatment following siKMT5C). Nevertheless, knockdown of KMT5C similarly increased transcript levels of multiple genes with the H3K9me3-/H4K20me3+ signature, paralleling the effects of A-196, with no additive or synergistic effect when both A-196 and KMT5C knockdown were combined (Figure 3G and Figure S4F).

**Figure 3.**
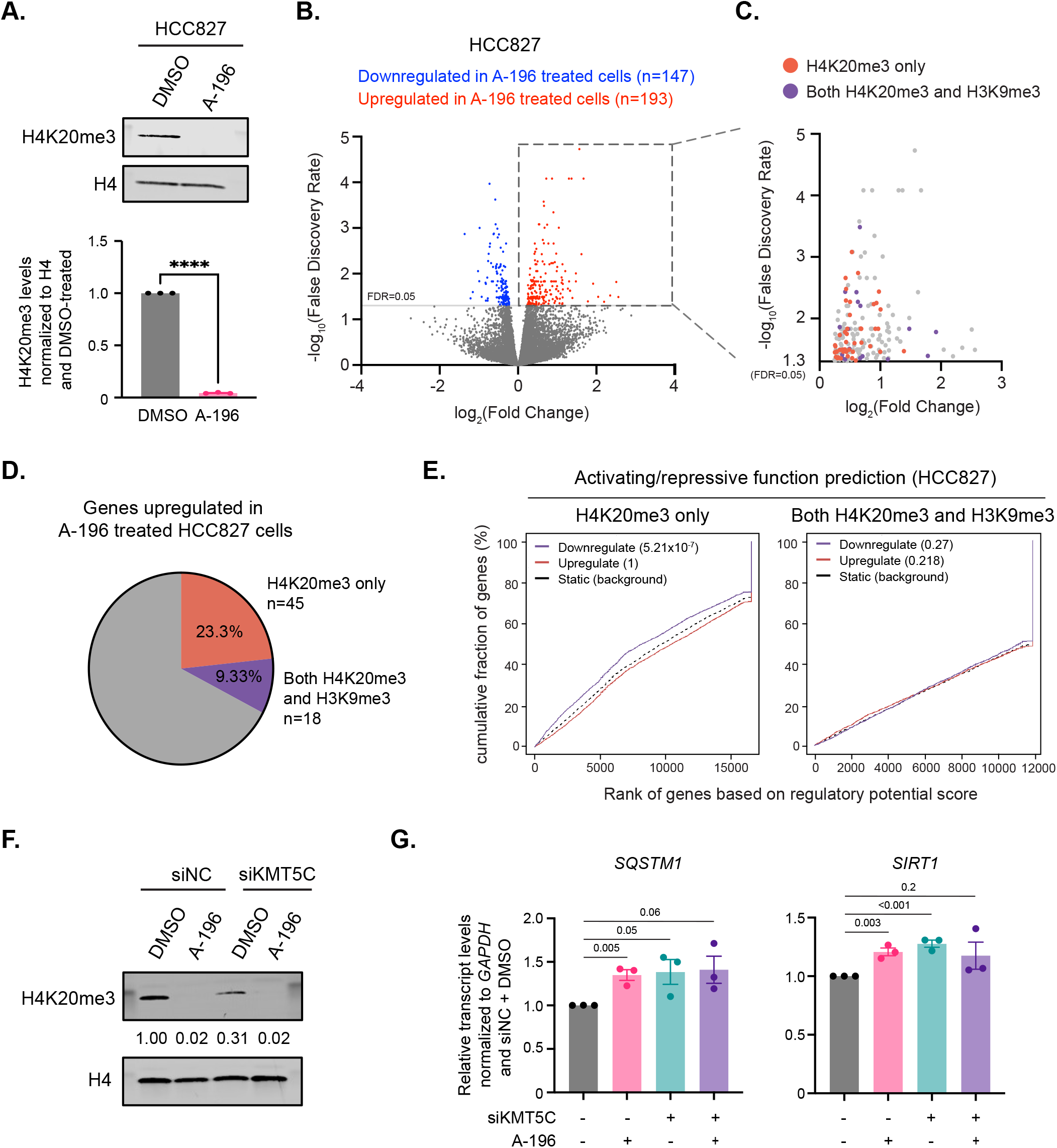
Genes with H3K9me3-/H4K20me3+ peaks in HCC827 cells exhibit more dynamic expression changes than those with H3K9me3+/H4K20me3+ peaks. **A**. Western blot analysis showing reduction of H4K20me3 following A-196 treatment in HCC827 cells. H4 was used as a loading control. Representative blot shown. n=3 biological replicates, quantified below. Data are presented as mean ± SEM, Welch’s t-test, ****P<0.0001 **B**. Volcano plot of differentially expressed genes after A-196 treatment in HCC827 cells. A cut-off of false discovery rate (FDR)=0.05 was used. **C**. Upregulated genes (FDR<0.05) in HCC827 cells annotated based on association with H3K9me3-/H4K20me3+ or H3K9me3+/H4K20me3+ peaks. **D**. Quantification of of upregulated genes (FDR<0.05) associated with each peak class in in HCC827 cells. **E**. BETA analysis predicting functional associations of H3K9me3-/H4K20me3+ and H3K9me3+/H4K20me3+ peaks in HCC827 cells. One sided KS-test was used to assess significance. **F**. Western blot analysis of H4K20me3 levels after KMT5C knockdown with or without A-196 treatment in HCC827 cells. H4 was used as a loading control. Representative blot shown; n=3 biological replicates. **G**. qRT-PCR analysis of transcript levels following KMT5C knockdown with or without A-196 treatment in HCC827 cells. Data were normalized to *GAPDH* and the siNC + DMSO negative control. n=3 biological replicates; two-sided Student’s t-test. P-values are indicated above bars represent.

RNA-seq analysis was also performed in PC9 cells treated with A-196 (Figure S5A). As in HCC827 cells, differentially expressed genes were enriched in EMT-related pathways (Figure S5B-D), and a greater fraction of these genes were associated with H3K9me3-/H4K20me3+ peaks compared to H3K9me3+/H4K20me3+ peaks (Figure S5E-F). However, BETA analysis in PC9 cells predicted no significant regulatory function for either group. We attribute this to the smaller number of differentially expressed genes following A-196 treatment in PC9 cells relative to HCC827 cells (Figure 3B and Figure S5B), which likely reflect lower baseline H4K20me3 levels in PC9 cells [5].

### KMT5C is recruited to H3K9me3-/H4K20me3+ peaks independent of HP1

H3K9me3 is generally regarded as an upstream modification of H4K20me3, reflecting the sequential deposition of H3K9me3 followed by H4K20me3 on chromatin. Canonically, KMT5C is recruited to H3K9me3-enriched regions through its interaction with HP1, which recognizes H3K9me3 via its chromodomain [12–14]. However, our data indicate that a subset of H4K20me3 peaks lacks H3K9me3 (Figure 1), suggesting that KMT5C may also be recruited to chromatin independent of its interaction with HP1.

KMT5C interacts with HP1 through its C-terminal domain, and deletion of this region (ΔCTD) abrogates this interaction [12,17,26]. To test whether HP1 binding is required for chromatin association, we generated HA-tagged constructs of full-length (FL) KMT5C and ΔCTD-KMT5C (Figure 4A). As expected, FL-KMT5C, but not ΔCTD-KMT5C, co-immunoprecipitated with HP1lil in HEK293T cells (Figure 4B), confirming loss of interaction.

**Figure 4.**
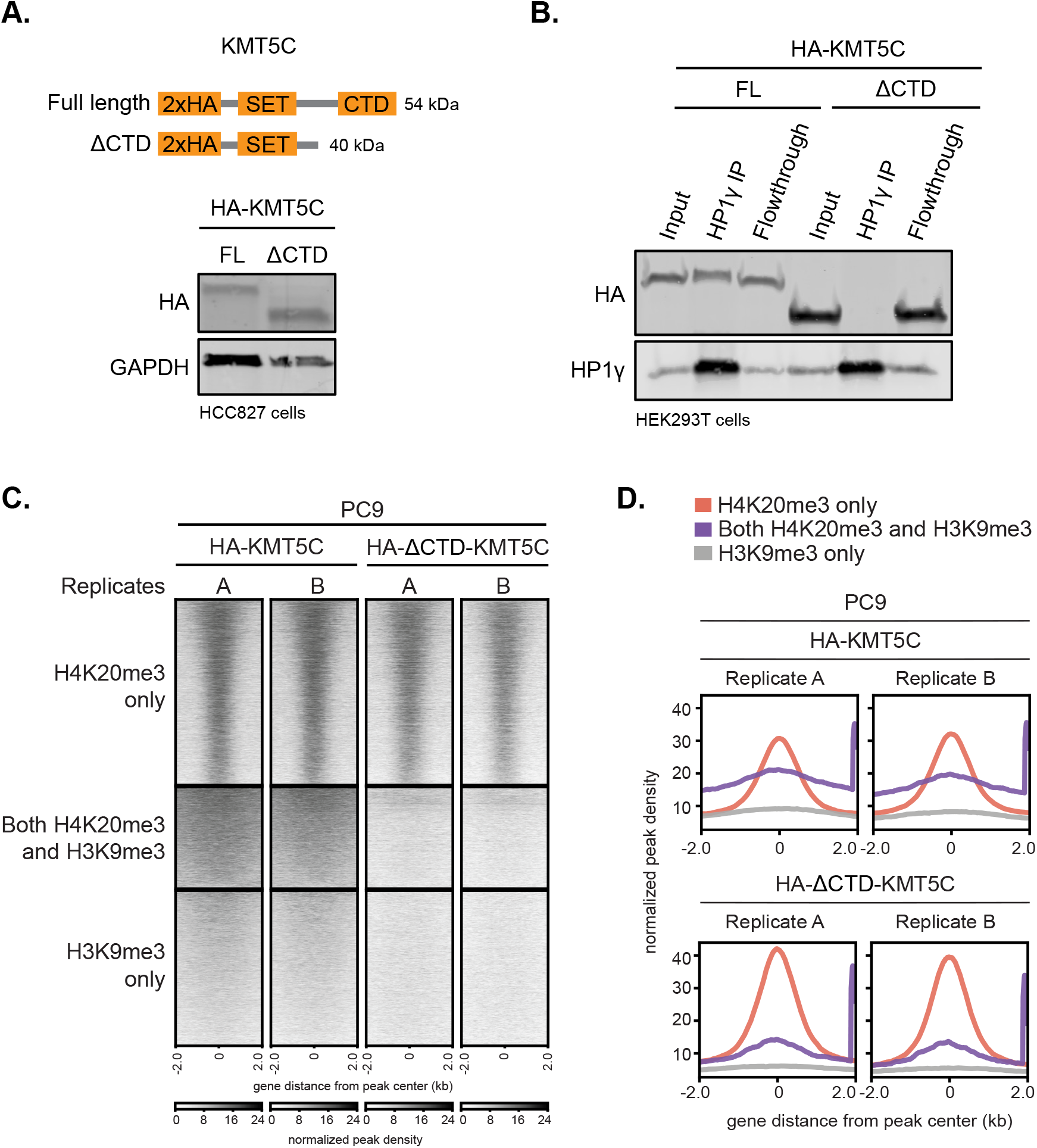
KMT5C is recruited to H3K9me3-H4K20me3 sites independent of its interaction with HP1. **A**. Schematics of generation of vectors that express HA-tagged full-length (FL) or C-terminal domain deletion mutant (ΔCTD) KMT5C. Vectors were validated by western blot using anti-HA antibody in HCC827 cells. GAPDH serves as a loading control. **B**. Co-immunoprecipitation analysis showing interaction between HA-FL-KMT5C and HP1L, which is lost for HA-ΔCTD-KMT5C mutant in HEK293T cells. Representative Western blot shown. n=3 biological replicates. **C**. Heatmaps of HA peaks clustered by H4K20me3 only, H3K9me3 only, or both H4K20me3 and H3K9me3 in PC9 cells transfected with HA-FL-KMT5C or HA-ΔCTD-KMT5C. **D**. Normalized peak density plots of HA peaks clustered by overlap with H4K20me3 only, H3K9me3 only, or both H4K20me3 and H3K9me3 peaks in PC9 cells transfected with HA-FL-KMT5C or HA-ΔCTD-KMT5C.

We next assessed whether the ΔCTD mutation altered chromatin binding. Overexpression of HA-tagged FL-or ΔCTD-KMT5C, followed by CUT&RUN using anti-HA revealed that both HA-KMT5C constructs were concentrated in more localized regions in the H4K20me3+ only subclass, similar to the observations for endogenous KMT5C (Figure 2B). Conversely, only the FL-KMT5C was enriched in the H3K9me3+/H4K20me3+ subclass, and the enrichment was distributed across the entire loci, similar to distribution of the endogenous KMT5C within this subclass (Figure 2A and Figure S6A). Unlike FL-KMT5C, ΔCTD-KMT5C was not enriched at H3K9me3+/H4K20me3+ peaks (Figure 4C and Figure S6B). Unexpectedly, ΔCTD-KMT5C remained localized at H3K9me3-/H4K20me3+ peaks at levels equal to or greater than FL-KMT5C (Figure 4D and Figure S6C). These findings demonstrate that KMT5C utilizes a mechanism other than an HP1-dependent mechanism to deposit H4K20me3 at H3K9me3-/H4K20me3+ regions.

### Identification of ZNF280C as a previously unknown interactor of KMT5C

To identify novel interactors of KMT5C that mediate HP1-independent recruitment to chromatin, we employed the proximity labeling technique BioID (Figure S7A). We used an HA-tagged BirA ligase enzyme (HA-BirA*) in three configurations: i) HA-BirA* alone, ii) HA-BirA* fused to GFP with a nuclear localization signal (HA-BirA*-GFP-NLS), and iii) HA-BirA* fused to KMT5C (HA-BirA*-KMT5C) (Figure 5A). The first two served as negative controls to distinguish nonspecific biotinylation events from those specific to KMT5C. We also generated HA-BirA*-ΔCTD-KMT5C to capture interactors of KMT5C independent of the HP1-binding domain. Importantly, HA-BirA*-ΔCTD-KMT5C retained proper nuclear localization (Figure 5B). To avoid streptavidin bead saturation from self-biotinylation, BirA*-containing vectors were depleted using an anti-HA antibody prior to streptavidin pull-down for proteomics analysis (Figure S7B). The sensitivity of the assay was confirmed by the enrichment of multiple HP1 isoforms exclusively in the FL-KMT5C group but not in ΔCTD-KMT5C samples (Figure 5C and Figure S7C).

**Figure 5.**
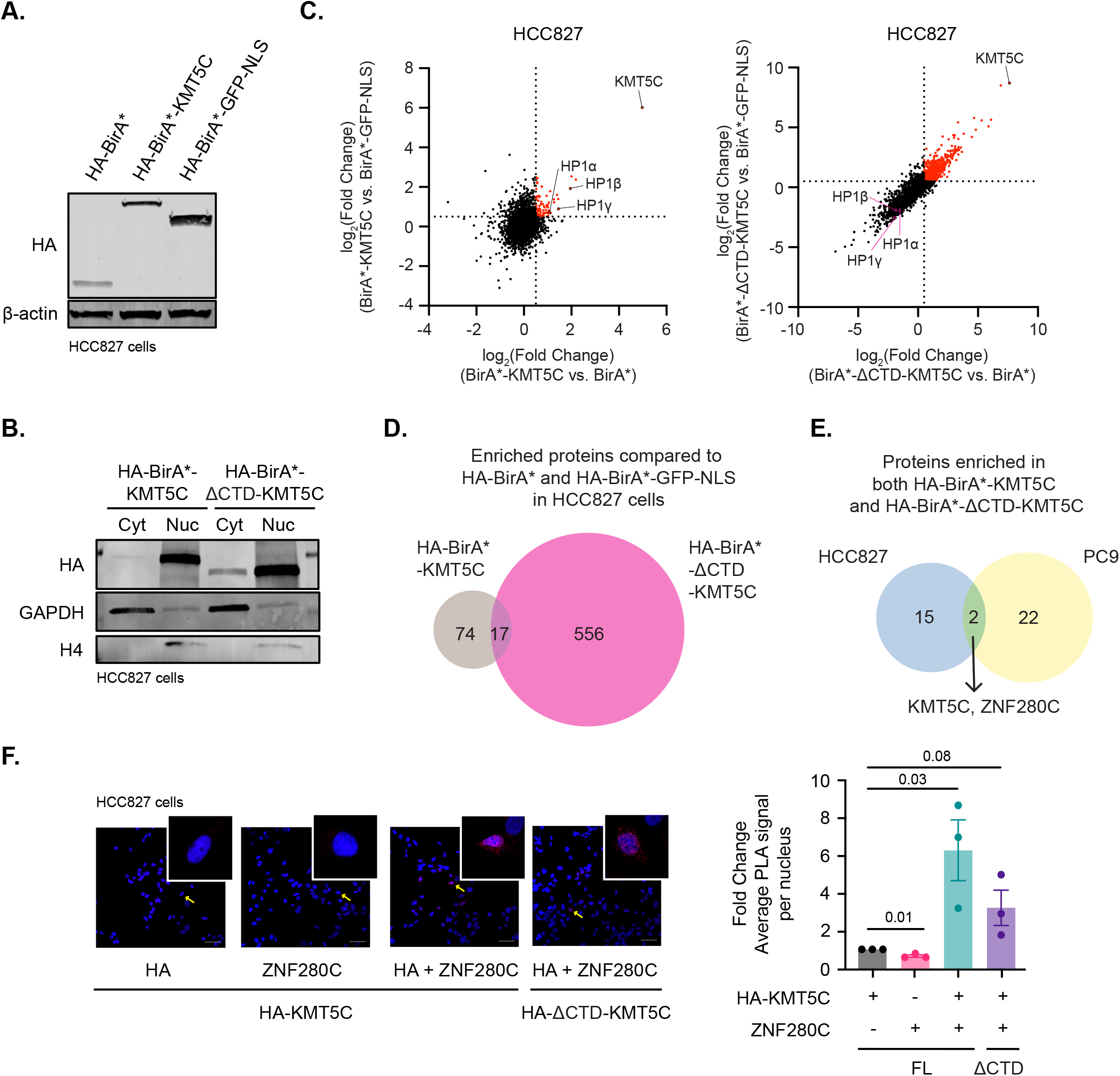
Identification of ZNF280C as a previously unknown interactor of KMT5C. **A**. Western blot validation of HA-BirA*, HA-BirA*-KMT5C, and HA-BirA*-GFP-NLS vectors. β-actin serves as a loading control. **B**. HA-BirA*-ΔCTD-KMT5C still retains the ability to be localized in the nucleus. GAPDH serves as a cytoplasmic marker and H4 as a nuclear marker. **C**. Proteomics analysis of enriched proteins in HA-BirA*-KMT5C or HA-BirA*-ΔCTD-KMT5C compared to either HA-BirA* or HA-BirA*-GFP-NLS in HCC827 cells. Cut-off of log_2_(fold change) of 0.5 was used. n=3 biological replicates. **D**. Identification of proteins that are enriched in both HA-BirA*-KMT5C FL and HA-BirA*-ΔCTD-KMT5C compared to either HA-BirA* or HA-BirA*-GFP-NLS in HCC827 cells. **E**. Identification of proteins enriched in HA-BirA*-KMT5C and HA-BirA*-ΔCTD-KMT5C in both HCC827 and PC9 cells. **F**. Proximity ligation assay (PLA) to validate interaction between KMT5C and ZNF280C in HCC827 cells. Cells that were probed with either HA or ZNF280C antibody serve as negative controls. Fold change of average PLA signal per nucleus is shown. n=3 biological replicates. Two-sided Student’s t-test was performed to evaluate statistical significance.

We next prioritized proteins enriched in both FL-KMT5C and ΔCTD-KMT5C datasets as candidate interactors at H3K9me3-/H4K20me3+ peaks (Figure 5D and Figure S7D). KMT5C and ZNF280C were the only proteins consistently enriched in both HA-BirA*-KMT5C and HA-BirA*-ΔCTD-KMT5C samples across both HCC827 and PC9 cell lines, identifying ZNF280C as a potential novel interactor of KMT5C that associates in an HP1-independent manner (Figure 5E). To validate this interaction, we performed a proximity ligation assay (PLA) in HCC827 cells, which detects proteins within ∼40 nm of one another [27]. A significantly higher PLA signal was observed in cells probed with both HA and ZNF280C antibodies compared to controls probed with either antibody alone. This enrichment was detected for both HA-FL-KMT5C and HA-ΔCTD-KMT5C, supporting that ZNF280C is in close proximity to KMT5C independent of HP1 binding (Figure 5F and Figure S7E).

### ZNF280C is involved in transcriptional regulation of genes associated with H3K9me3-/H4K20me3+ peaks

Having identified ZNF280C as a novel interacting partner of ΔCTD-KMT5C (Figure 5), we hypothesized that ZNF280C would localize specifically to H3K9me3-/H4K20me3+ sites, but not to H3K9me3+/H4K20me3+ peaks. To test this, we analyzed publicly available ZNF280C ChIP-seq data (GSE181855) and compared it with our H4K20me3 CUT&RUN dataset [28]. Although the ChIP-seq was performed in a different cancer model (RKO colorectal cancer cells), ZNF280C peaks were specifically enriched at H3K9me3-/H4K20me3+ regions, but not at H3K9me3+/H4K20me3+ peaks (Figure 6A-B). Consistent with this, TCGA panCancer Atlas data revealed a positive correlation between *ZNF280C* and *KMT5C* mRNA levels (Figure 6C). The data were encouraging considering the correlation was only slightly lower than that observed between the bonafide target of KMT5C, *CBX3(HP1⍰)* and *KMT5C* (Figure 6C). To directly test whether ZNF280C contributes to transcriptional regulation of genes with H3K9me3-/H4K20me3+, siRNA-mediated knockdown of *ZNF280C* was conducted. Using various siRNAs targeting *ZNF280C*, knockdown resulted in increased expression of *SIRT1* and *SQSTM1* (Figure 6D), two genes we previously identified as H3K9me3-/H4K20me3+ targets under KMT5C regulation (Figure 3G).

**Figure 6.**
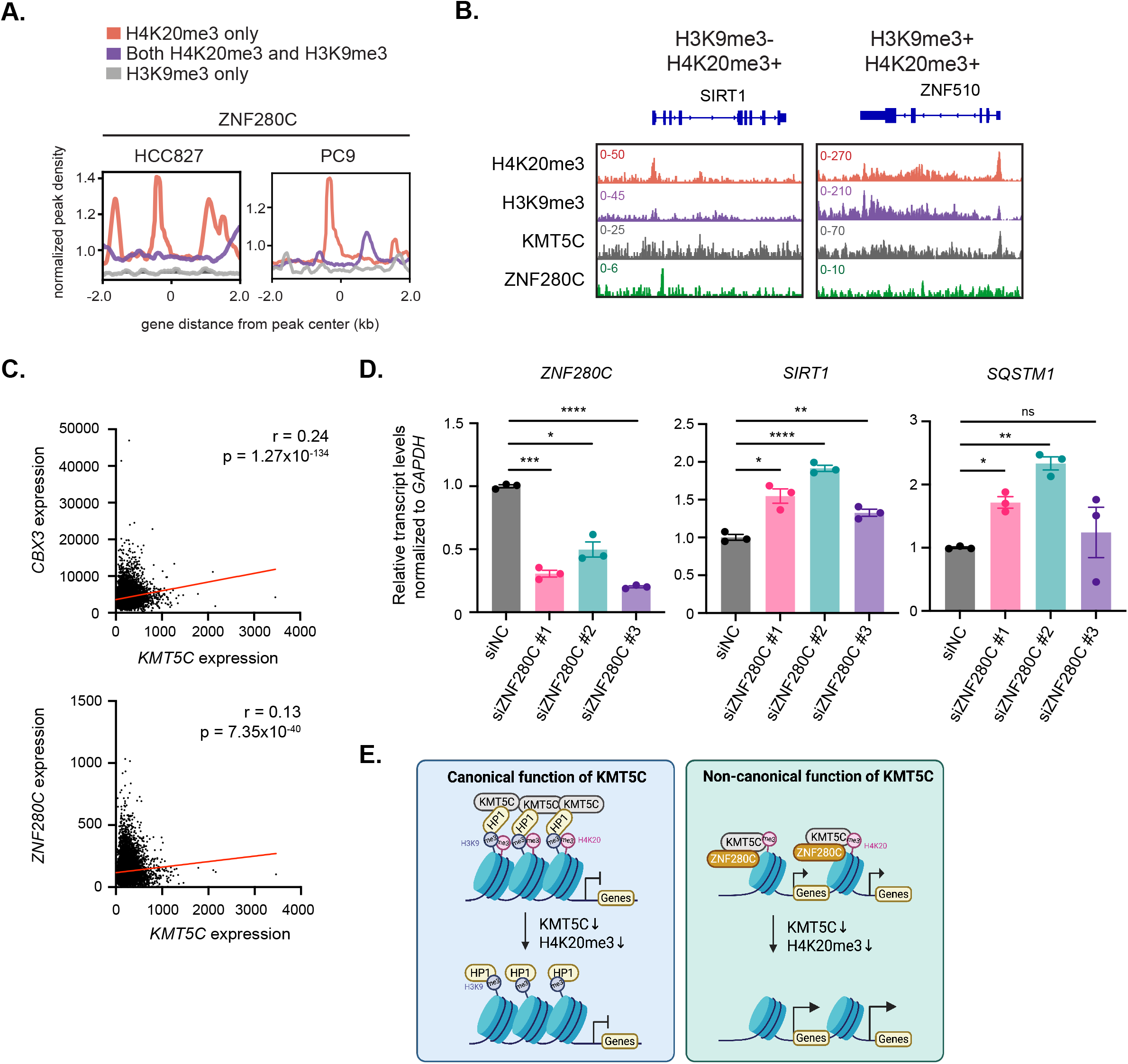
ZNF280C is involved in transcriptional regulation of H3K9me3-/H4K20me3+ genes. **A**. Normalized peak density plots to evaluate ZNF280C binding on HCC827 H3K9me3-/H4K20me3+ or H3K9me3+/H4K20me3+ peaks. ZNF280C ChIP-seq data was acquired from GSE181855. **B**. IGV view of representative genes with H3K9me3-/H4K20me3+ peaks (SIRT1) and H3K9me3+/H4K20me3+ peaks (ZNF510) with KMT5C and ZNF280C peaks. One of two (n=2) biological replicates of HCC827 CUT&RUN peaks is shown. **C**. Correlation between mRNA levels of *KMT5C* and *ZNF280C*, as well as that between mRNA levels of *KMT5C* and *CBX3*. Data was acquired from cBioPortal using TCGA panCancer atlas. Spearman test was used to calculate Spearman correlation value (r) and statistical significance. **D**. qRT-PCR analysis of transcript levels after ZNF280C knockdown. Data were normalized to *GAPDH* and shown relative to siNC. Technical replicates of one of three (n=3) biological replicates are shown. Two-sided Student’s t-test was performed to evaluate statistical significance. *P<0.05, **P<0.01, ***P<0.001, ****P<0.0001. **E**. Working model depicting the collaboration between KMT5C and ZNF280C in non-canonical deposition of H4K20me3 independent of H3K9me3.

Together, these findings indicate that ZNF280C likely collaborates with KMT5C at H4K20me3-marked, H3K9me3-depleted regions, revealing a non-canonical function of KMT5C that helps safeguard against excessive gene expression (Figure 6E).

## DISCUSSION

Our findings uncover a previously unrecognized subclass of H4K20me3-marked chromatin that lacks H3K9me3, a subclass that appears to be established through a non-canonical mechanism. Historically, H4K20me3 has been viewed as a constitutive repressive mark at repetitive regions of the genome, deposited downstream of H3K9me3 and HP1 recruitment [12–14]. However, our data challenge this model, showing that a substantial portion of H4K20me3 peaks exist independently of H3K9me3. While it is plausible that this subset of H4K20me3 peaks is deposited downstream of H3K9me3, with subsequent removal of H3K9me3, our CTD mutant suggest otherwise. Our data suggests that this newly identified H4K20me3 peaks does not follow the conventional route of deposition. These peaks are enriched at transcription start sites and co-localize with active histone modifications. In support of the activating marks, genes closest to this subclass are correlated with the highest gene expression profiles in comparison to all other marks evaluated. These observations reveal unexpected functional and mechanistic diversity in H4K20me3 biology.

The identification of the H3K9me3-/H4K20me3+ signature at promoters of genes involved in replication, RNA processing, and cell cycle regulation (Figure S1B, S4A, S5) suggests that this signature participates in fine-tuning expression. Indeed, inhibition or depletion of KMT5C led to enhanced transcriptional activation of some of the genes involved in these pathways (Figure 3, S4, S5), implying that H4K20me3 may act as a molecular “brake” to prevent excessive overexpression in active chromatin contexts. This concept extends upon emerging evidence that repressive marks can function in a modulatory capacity, buffering transcriptional output to maintain cellular homeostasis. Of particular interest is a clear division in presence or absence of H3K4me3 marks within the H4K20me3 subpopulation (Figure S6A), with about 1/3 of the loci having the H3K4me3 activation mark and 2/3 devoid of this mark. The functional differences between these unique and distinct subpopulations will be of interest to dissect in future studies. Mechanistically, our work establishes that KMT5C recruitment to these sites does not require HP1 (Figure 4, S6). Instead, the enzyme can localize to H3K9me3-depleted chromatin through an alternative pathway that may involve the zinc finger protein ZNF280C (Figure 5). ZNF280C emerged from our BioID screen as a consistent interactor of both full-length and ΔCTD-KMT5C, and its knockdown phenocopied KMT5C loss by derepressing a subset of H3K9me3-/H4K20me3+ associated genes. These results identify ZNF280C as a potential adaptor that may help to guide KMT5C to specific genomic loci independent of H3K9me3, thereby expanding the known repertoire of mechanisms through which histone methyltransferases are targeted to chromatin, albeit studies will need to be undertaken to verify these findings and to determine if the interaction is direct or indirect.

Functionally, the existence of dual H4K20me3 subclasses, one canonical and HP1-dependent, the other non-canonical and potentially ZNF280C-dependent, suggests a model in which KMT5C integrates signals from both constitutive heterochromatin and more dynamic euchromatic regions. This flexibility may be particularly relevant in cancer, where epigenetic rewiring enables transcriptional plasticity and drug resistance. Our prior work linking H4K20me3 to EGFR inhibitor resistance in lung cancer supports the possibility that non-canonical H4K20me3 contributes to adaptive transcriptional programs [5].

Future studies should validate and address how ZNF280C recognizes its target loci, whether additional cofactors modulate KMT5C recruitment, and how these mechanisms are influenced by cellular context or stress. Together, our results redefine the relationship between H4K20me3 and H3K9me3, uncovering an unanticipated layer of chromatin regulation that operates outside classical heterochromatin to maintain transcriptional balance in cancer cells.

## MATERIALS & METHODS

### Cell culture

HCC827 and PC9 cells were cultured in RPMI-1640 media (Cytiva #SH30027.FS) with 10% fetal bovine serum (FBS, BioTechne #S11150) and 1% penicillin/streptomycin (Pen/Strep, Cytiva #SV30010). HEK293T cells were cultured in DMEM media with 10% FBS and 1% Pen/Strep. All cell lines were regularly tested for mycoplasma by MycoAlert Mycoplasma Detection Kit (Lonza #LT07-418).

For BioID, HCC827 and PC9 cells that stably express a TetR repressor (Addgene #17492) were used. HEK293T cells were transfected with lentiviral constructs (psPAX2 Addgene #12260 and pMD2.G Addgene #12259) and TetR-expressing plasmid using lipofectamine 2000 (Invitrogen # 11668019). Viral supernatant was collected 48 hours post transfection. Viral particles were collected by centrifugation and used for transduction with 2µg/mL polybrene (Sigma Aldrich # TR1003G). Transduced cells were selected using 5µg/mL blasticidin (Gibco # A1113903) and single cell dilution was performed to isolate single cell clones.

### Plasmids

To generate HA-KMT5C, KMT5C was cloned into pMH-HA (Addgene #101767) using Gateway cloning (Invitrogen #11791020). For HA-ΔCTD-KMT5C, CTD (Δ347-462) was deleted using QuikChange Lightning Site-Directed Mutagenesis Kit (Agilent Technologies #210518) and the primers (5’-3’) GGGCCCCAGTGCTCTGAGACCCAGCTTT and AAAGCTGGGTCTCAGAGCACTGGGGCCC. For BioID, KMT5C was cloned into HA-BirA*-pDEST-N-pcDNA5/FRT/TO (Addgene #118375) using Gateway cloning (Invitrogen #11791020).). For HA-BirA*-ΔCTD-KMT5C, QuikChange Lightning Site-Directed Mutagenesis Kit (Agilent Technologies #210518) was used using the same primers as used for HA-ΔCTD-KMT5C. For HA-BirA*-GFP-NLS, GFP-NLS was cloned from pEGFP-C1 EGFP-3XNLS (Addgene #58468) and inserted into HA-BirA*-pDEST-N-pcDNA5/FRT/TO with BstB1 and Apa1 (New England Biolabs #R0519 and #R0114).

### CUT&RUN

CUT&RUN was performed using the EpiCypher® CUTANA™ CUT&RUN Protocol v2.0 as previously described [30]. pAG-MNase was a gift from the laboratory of Patrick Murphy (University of Rochester). Briefly, 5*10^5^ cells were incubated with Concanavalin-A beads (Bangs Laboratories, #BP531) followed with an overnight incubation with primary antibody at 4°C. Samples were then incubated for 10 min with pAG-MNase at room temperature followed by a 2hr 4°C incubation with 100mM CaCl_2_ to activate cleavage. MNase activity was halted with Stop Buffer and incubated for 10 min at 37°C. DNA was purified using the Qiagen MinElute Kit (Qiagen, #28004). Reactions were performed in duplicate and library preparation was performed using Illumina adapters and the NEBNext Ultra II DNA Library Kit for Illumina (NEB, #E7645). Libraries were pooled together and sequenced on the Illumina NovaSeq X platform (150bp PE) by the University of Rochester Genomics Research Center. For CUT&RUN, 0.5µg of the following primary antibodies was used: H4K20me3 (Abcam, #ab9053), H3K9me3 (ActiveMotif, #39162), H3K27me3 (Cell Signaling, #9733), H3K27ac (Abcam, #ab4729), H3K4me3 (Epicypher, #13-0041), H4K20me1 (Abcam, #ab9051), KMT5C (Sigma, #HPA052294), HA Tag (Epicypher, #13-2010), and IgG (Epicypher, #13-0042).

CUT&Run, and public CUT&Tag raw sequencing data were subjected to quality trimming using fastp (v.0.23.1) and read alignment to the GRChg38 reference genome (NCBI assembly GCF_000001405.4) using Bowtie2 (v.2.3.5.1). Mapped reads with mapping quality < 10 were removed. Duplicate aligned reads were marked using picard (v.2.12.0). Reads mapped to unplaced scaffolds, the Y chromosome, and blacklisted regions (The ENCODE Blacklist, hg38-blacklist.v2.bed, https://github.com/Boyle-Lab/Blacklist/tree/master) were also excluded. BAM files were converted to bedGraph format using bedtools (v.2.30.0) and subjected to SEACR (v.1.3) to identify enriched regions by selecting the top 1% of regions by AUC. RPKM-normalized BigWig files were created using DeepTools bamCoverage (v3.5.1) with the following parameters: --normalizeUsingRPKM --ignoreForNormalization chrX --extendReads --binSize10. Co-localization of peaks of H4K20me3 and H3K9me3 was using bedtools and the heat maps and metaplots of the corresponding regions were plotted with Deeptools. Customized R scripts were used to annotate peaks to the closest TSS with TxDb.Hsapens.UCSC.hg38.refGene (v.3.15) annotation.

For CUT&RUN-qPCR, CUT&RUN was performed using EpiCypher® CUTANA™ CUT&RUN kit (Epicypher #14-1048). Input genomic DNA was isolated from 5*10^5^ cells using PureLink™ Genomic DNA Mini Kit (Invitrogen #K182001). Following CUT&RUN, qPCR was performed using QuantiNova SYBR Green PCR Kit (Qiagen #208056) with the following primers.

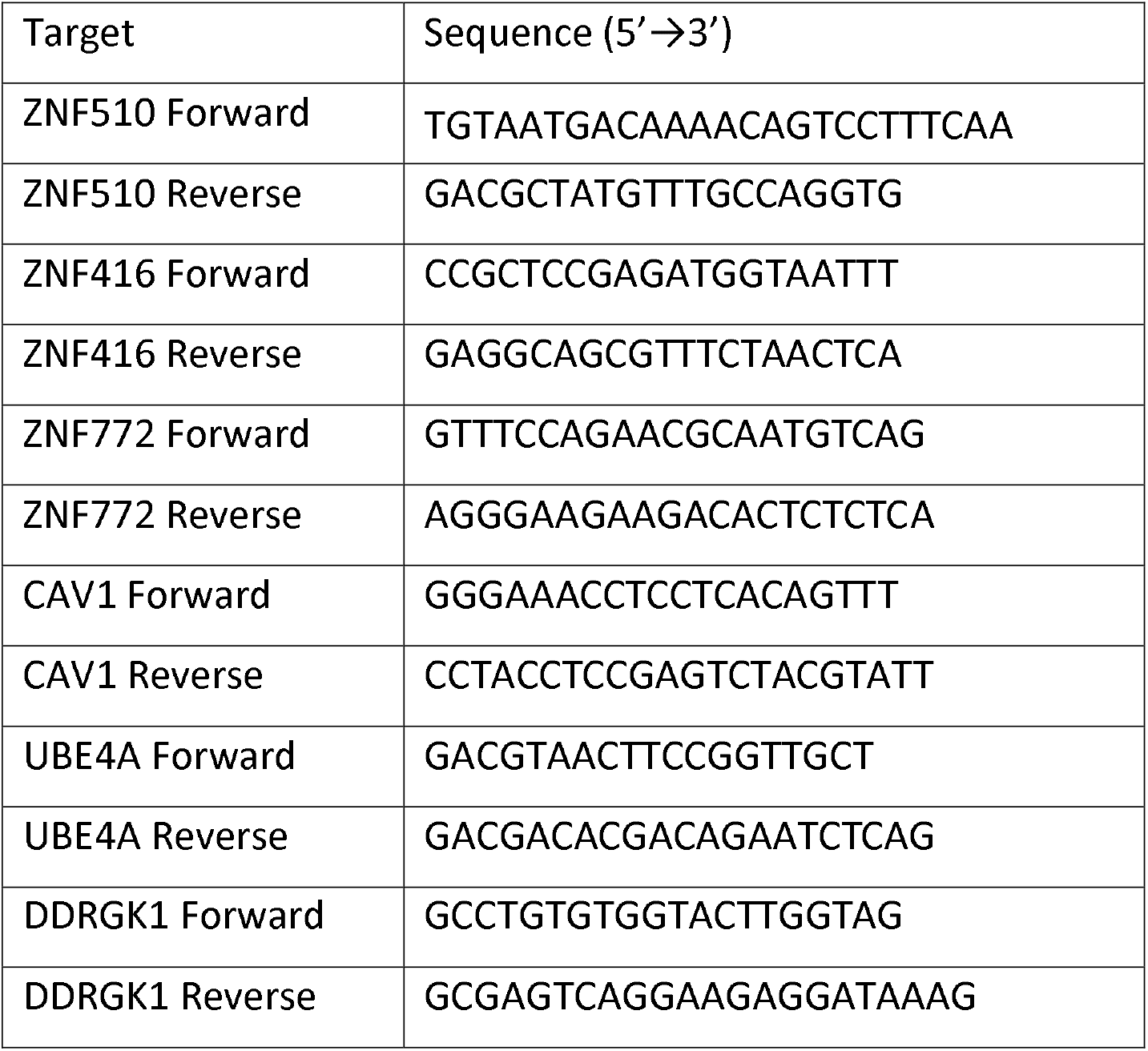

### Protein extractions and immunoblotting

For whole-cell lysates, cells were lysed using RIPA buffer (150mM NaCl, 50mM Tris-HCl, 1% NP-40, 0.5% sodium deoxycholate, and 0.1% SDS) supplemented with 1x protease inhibitor (Thermo Scientific #A32955). For nuclear fractionation, previously reported methods from Gagnon, K. et al [31] and Zhu, C. et al [32] were used. Total protein amounts were quantified using Pierce™ BCA Protein Assay Kit (Thermo Scientific #3225).

For immunoprecipitation, cells were lysed using IP lysis buffer (50mM Tris-Cl, 150mM NaCl, 1mM EDTA, and 1% TritonX-100) supplemented with 1x protease inhibitor (Thermo Scientific #A32955). Lysates were pre-cleared using dynabeads G (Invitrogen #10004D) for 2 hours on a rotator at 4°C. An aliquot was stored as input for western blot analysis. Lysates were incubated with HP1⍰ (Proteintech #11650-2-AP, 1.5µg per 2mg lysates) overnight on a rotator at 4°C. Next day, Dynabeads G were added and incubated on a rotator at 4°C for 1 hour. Bead/antibody complex was washed four times with IP lysis buffer and proteins were eluted with 4x SDS loading buffer (Licor #928-40004) by boiling at 95°C for 10 minutes.

Histones were extracted using previously reported method from Shechter, D. et al [33]. Proteins were resolved using 4–20% criterion™ TGX™ precast midi protein gel (Biorad #5671093 or #5671094), transferred to Immobilon™-FL PVDF membranes (Sigma-Aldrich #IPFL00005), blocked for 1 hour at room temperature using Intercept® (PBS) Blocking Buffer (Licor #927-70003), and probed with primary antibodies at 4°C overnight. The following antibodies were used: H4K20me3 (Active Motif #39671, 1:500), H4 (Active Motif #61299, 1:500), HA (Cell Signaling Technology #2367S, 1:1000), HA (Invitrogen #26183, 0.2µg/mL), HA (Cell Signaling Technology #3724S, 1:1000), GAPDH (Cell Signaling Technology #2118S, 1:1000), HP1⍰ (Proteintech #11650-2-AP, 1:1000), β-actin (Cell Signaling Technology #3700S, 1:1000). Immunoblots were incubated with IRDye secondary antibodies (Licor #925-32211, #925-68071, #925-32210, or #925-68070, Invitrogen #A27042) for 1 hour at room temperature and imaged with the Licor Odyssey CLX.

### siRNA knockdown

Cells were transfected with siRNAs using Lipofectamine RNAiMAX (Invitrogen #13778150). 4 hours after transfection, media was replaced with complete media. Cells were collected 72 hours (ZNF280C) or 120 hours (KMT5C) post-transfection. The following siRNAs were used: KMT5C (Dharmacon #J-018622-20-0005, 10nM), ZNF280C #1 (Thermo Fisher Silencer® Select #s31065, 10nM), ZNF280C #2 (Thermo Fisher Silencer® Select #s31066, 10nM), and ZNF280C #3 (Thermo Fisher Silencer® Select #s31067, 10nM).

### mRNA qPCR and RNA-seq

Total RNA was extracted from cells using miRNeasy Micro Kit (Qiagen #217004). Equal amounts of RNA were used to synthesize cDNA using SuperScript™ IV VILO™ Master Mix (Invitrogen #11756050). qPCR was performed using TaqMan™ Fast Advanced Master Mix for qPCR (Applied Biosystems #4444963) with the following primers: KMT5C (Applied Biosystems, #Hs00261961_m1), SIRT1 (Applied Biosystems, # Hs01009006_m1), SQSTM1 (Applied Biosystems, #Hs01061917_g1), ZNF280C (Applied Biosystems, # Hs01585681_m1), and GAPDH (Applied Biosystems, #Hs99999905_m1).

For RNA-seq, total RNA was sent to Novogene for QC, library construction and sequencing. RNA samples were sequenced using Illumina platform following paired-end sequencing protocol with read length 150 bp and at least 40 million paired-end reads per sample. Data quality control (adapter removal, phred quality cutoff >=Q30, minimum read length 50 bp after quality filtering) was performed using fastp (version 0.23.2) tools [34]. Reads were aligned to human (GRCh38) reference genome using STAR aligner (version 2.7.11b)[35]. Strand specificity was determined with RSeQC package (version 4.0.0) [36] followed by counting the reads mapped to each gene in each sample with FeatureCounts (version 2.0.1) [37] and raw counts matrix was generated. Differential Expression (DE) analysis was performed using edgeR Bioconductor package (version 4.0.16) [38], and the P-values for multiple testing were adjusted using Benjamini and Hochberg method [39]. FDR < 0.05 was chosen as a cutoff for the significant differential expression. Gene Set Enrichment Analysis (GSEA) was performed with GSEA software (version 4.4.0) [40,41] and Hallmark gene set from MSigDB collection [42]. Transcript Per Million (TPM) counts were generated using TPMCalculator tool (version 0.0.4) and used to generate any heatmaps. Custom visualizations were created using R-packages ggplot2 (version 3.5.1) [43], EnhancedVolcano (version 1.20.0) [44] and ComplexHeatmap (version 2.18.0) [45].

### BioID

HCC827 and PC9 cells that stably express a TetR repressor were used. These cells were grown in 5ug/mL blasticidin RPMI media supplemented with 10% FBS and 1% pen/strep. For BioID, cells were seeded in 15 cm plates in DMEM media (Gibco #11965118) supplemented with 10% FBS and 1% pen/strep such that cells reach 70∼90% confluency next day. DMEM media was used for BioID as DMEM does not contain biotin. Next day, cells were transfected with HA-BirA*, HA-BirA*-GFP-NLS, HA-BirA*-KMT5C, or HA-BirA*-ΔCTD-KMT5C using lipofectamine 3000 (Invitrogen #L3000015). 4 hours after transfection, media was replaced with DMEM media containing 2 µg/mL doxycycline (Fisher Scientific #BP26531). 24 hours post transfection, cells were treated with 50 µM biotin (Thermo Scientific # 29129). Cells treated with biotin for 24 hours were collected and fractionated into cytoplasm and nuclear fractions. Pre-cleared nuclear lysates were first incubated with anti-HA antibody (Abcam #ab9110, 1µg per 200µg input proteins) to deplete nuclear lysates of HA-tagged proteins. Next day, antibody/protein complexes were separated using Dynabeads A (Invitrogen #10002D) on a rotator for 2 hours at 4°C. Flowthrough was subsequently used as an input for enrichment of biotinylated proteins using streptavidin beads (Thermo Scientific #88816) on a rotator overnight at 4°C. After overnight incubation, 80% of the beads were flash-frozen for mass spectrometry analysis. Remaining 20% of the beads were used for validation by western blot. Proteins were eluted from the beads with 4x protein loading buffer supplemented with 2mM biotin and 20mM DTT by boiling at 95°C for 10 minutes. Proteomics analysis was performed by IDeA National Resource for Quantitative Proteomics at the University of Arkansas Medical School. Differential expression analysis was performed using DEP R package (version 1.28.0) [46].

### Proximity Ligation Assay

2.5*10^5^ HCC827 cells were seeded in a 6-well plate. Next day, cells were transfected with HA-KMT5C or HA-ΔCTD-KMT5C using lipofectamine 3000 (Invitrogen #L3000015). The following day, cells were collected and 1*10^5^ cells were re-seeded in 8-well Chambered Cell Culture Slides (MatTek Corporation #CCS8). Next day, cells were fixed using 4% paraformaldehyde and permeabilized using 0.1% Triton-X100. Proximity ligation assay (PLA) was performed using Duolink® In Situ Red Starter Kit Mouse/Rabbit (Sigma-Aldrich #DUO92101-1KT) following manufacturer’s instruction. The following primary antibodies were used in the PLA: HA (Invitrogen #26183, 2µg/mL) and ZNF280C (Sigma #HPA051494, 2µg/mL).

## Supporting information

Supp BioID

## Data availability

Proteomics data are available in Supplementary File. All other raw data are available upon request from the corresponding author.

### Statistical analysis

All experiments were performed using 2 biological replicates for CUT&RUN, and 3 biological replicates for all other experiments. Statistical analysis was performed using Prism (GraphPad Version 10.5.0). Statistical tests used to determine the statistical significance of the difference between groups are specified in the figure legends.

## Author contributions

J.S. and A.L.K. conceived and designed the study. J.S. performed in vitro experiments and bioinformatic analysis for CUT&RUN, RNA-seq, and proteomics data. C.S. performed bioinformatic analysis for the CUT&RUN data. N.M. performed bioinformatic analysis for the CUT&RUN and RNA-seq data. C.D. performed CUT&RUN experiments. S.M.U. performed RNA-seq analysis. A.G. assisted in generating vectors for BioID. P.M.V. guided the CUT&RUN experiments and analysis of the data. J.S. and A.L.K. wrote and edited the manuscript.

## Acknowledgments

This study was funded in part by the National Institutes of Health (7R01CA205420 and 7R01CA226259) to A.L.K., (5R01CA250531) to P.M.V, and a grant from the NIH to the Purdue Institute for Cancer Research (44P30CA023168). Additional support was obtained from the Genomics Shared Resource at the Wilmot Cancer Institute supported in part by the University of Rochester Wilmot Cancer Institute Support Grant (1P30CA272302). J.S. was supported by a Ross Fellowship and Bilsland Fellowship from Purdue University. J.S. was also a recipient of Biomedical Science Scholarship from Asan Foundation. A.G. was supported by a Summer Research Fellowship from the Purdue Institute for Cancer Research.

