## Supplementary material for "KMT5C-H4K20me3 drives changes in epigenetic landscape independent of H3K9me3": Supp Figures

**Figure S1. A subset of H4K20me3 is deposited independent of H3K9me3, related to Figure 1.**

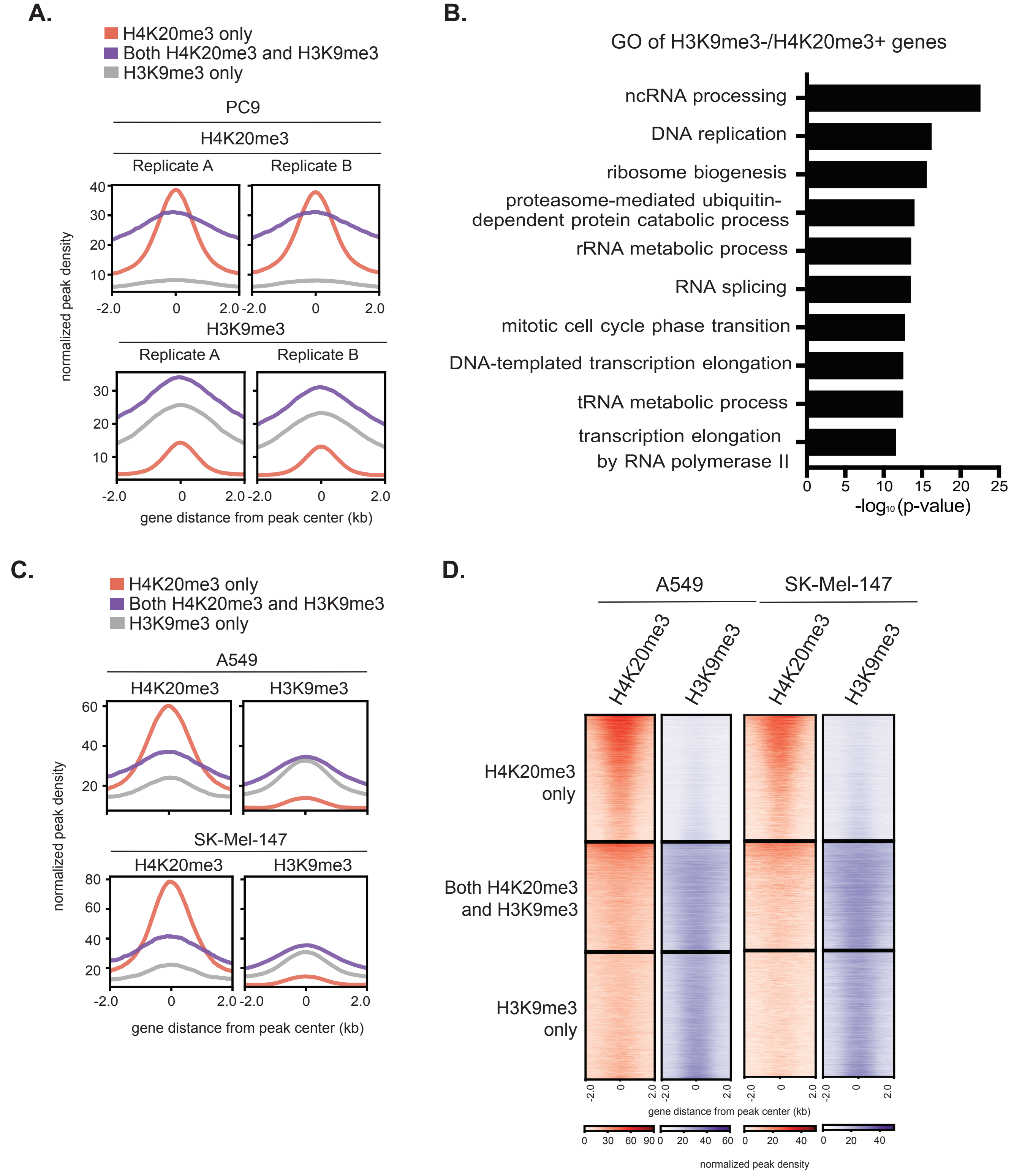

**Figure S2. H3K9me3-/H4K20me3+ peaks lack canonical repressive epigenetic signatures in PC9 cells, related to Figure 2.**

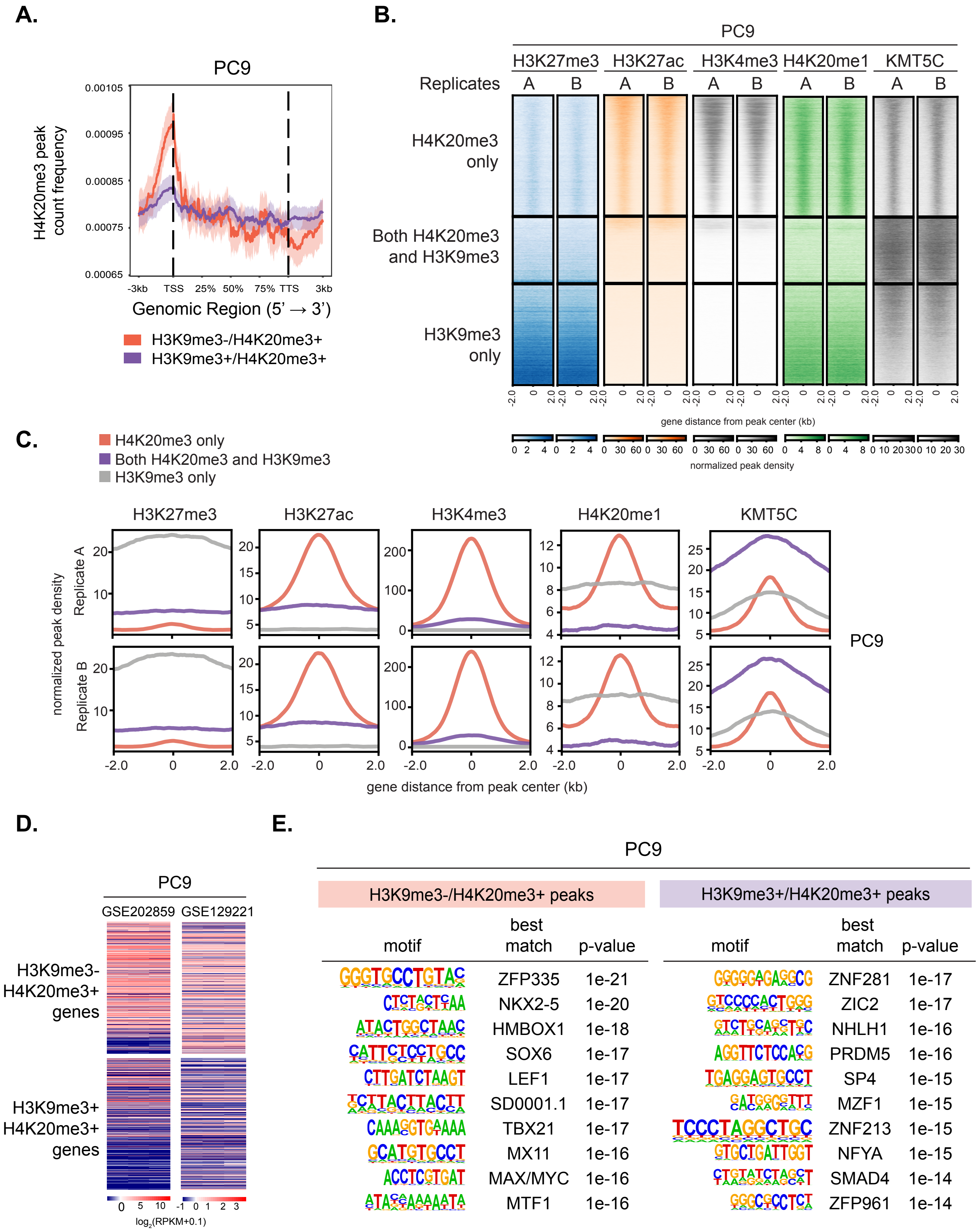

**Figure S3. H3K9me3-/H4K20me3+ peaks are related with highly-expressed genes in HCC827 cells, related to Figure 2.**

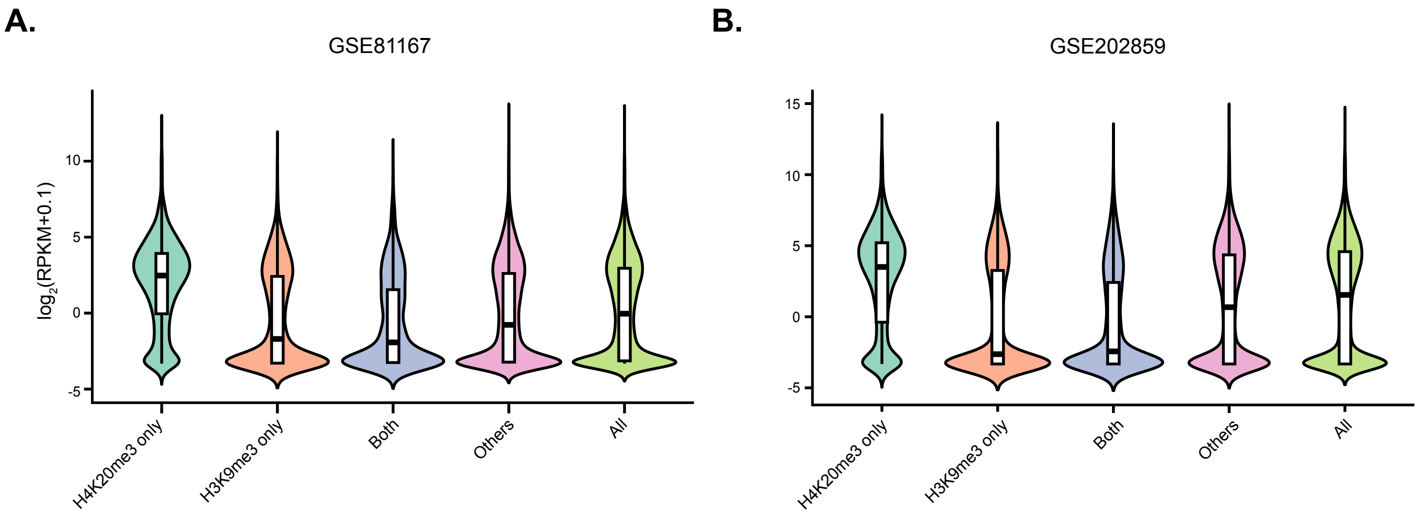

**Figure S4. Genes with H3K9me3-/H4K20me3+ peaks in HCC827 cells exhibit dynamic expression changes, related to Figure 3.**

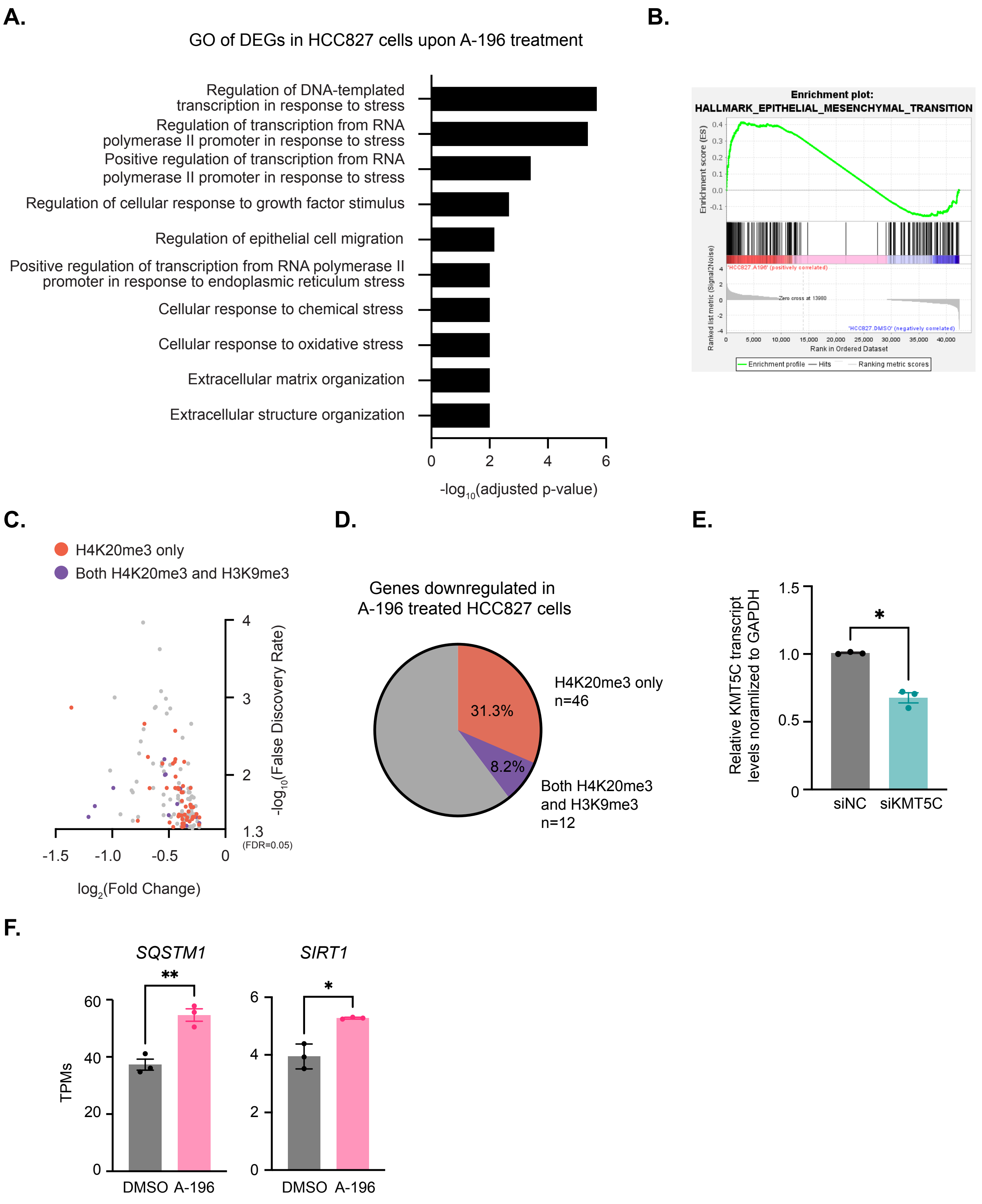

**Figure S5. Genes with H3K9me3-/H4K20me3+ peaks in PC9 cells are enriched for the epithelial to mesenchymal transition pathway and exhibit more dynamic expression changes than those with H3K9me3+/H4K20me3+ peaks, related to Figure 3.**

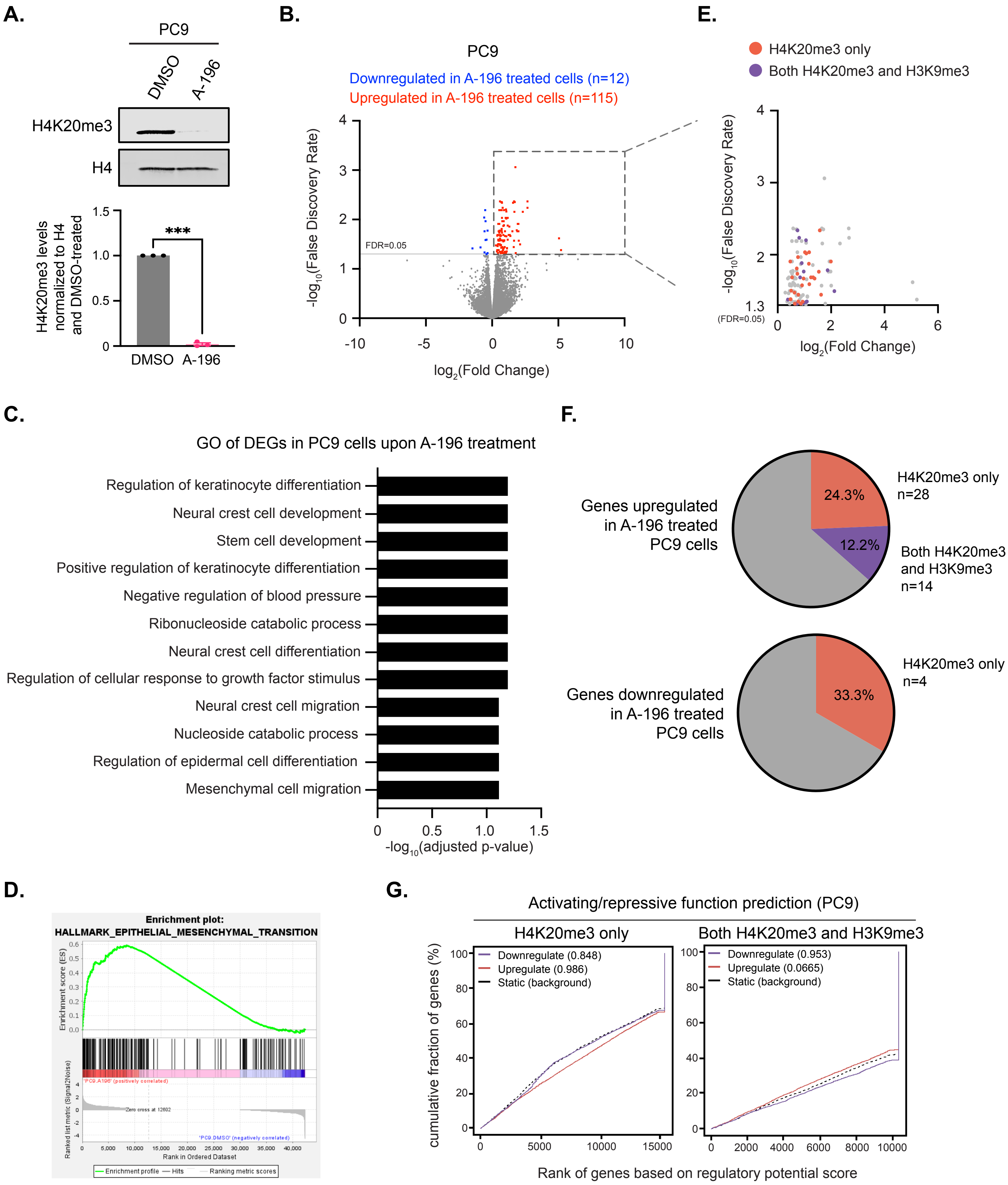

**Figure S6. HA-ΔCTD-KMT5C retains the ability to be recruited to H3K9me3-/H4K20me3+ sites, related to Figure 4.**

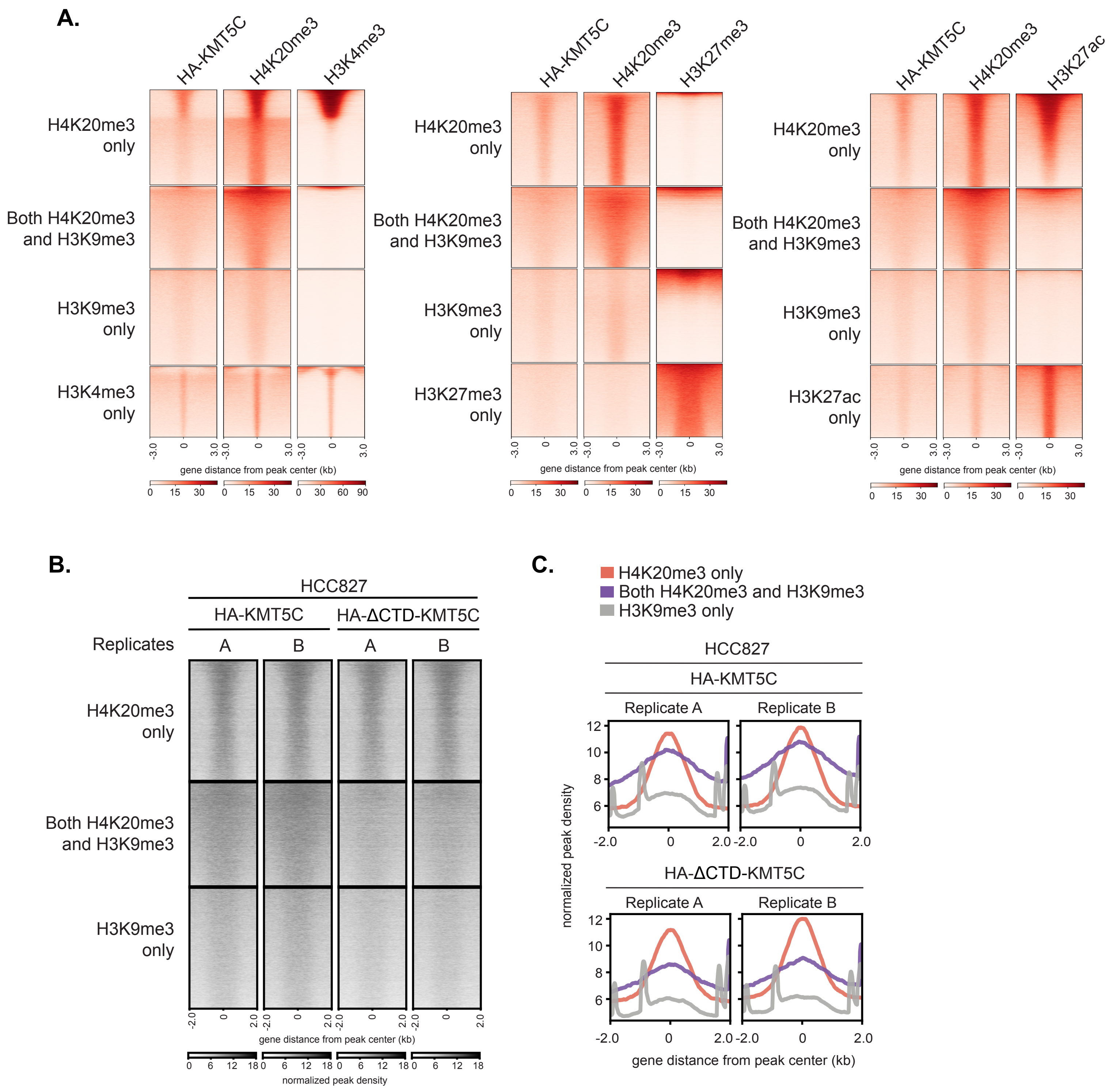

**Figure S7. Identification of novel KMT5C interactome using BioID, related to Figure 5.**

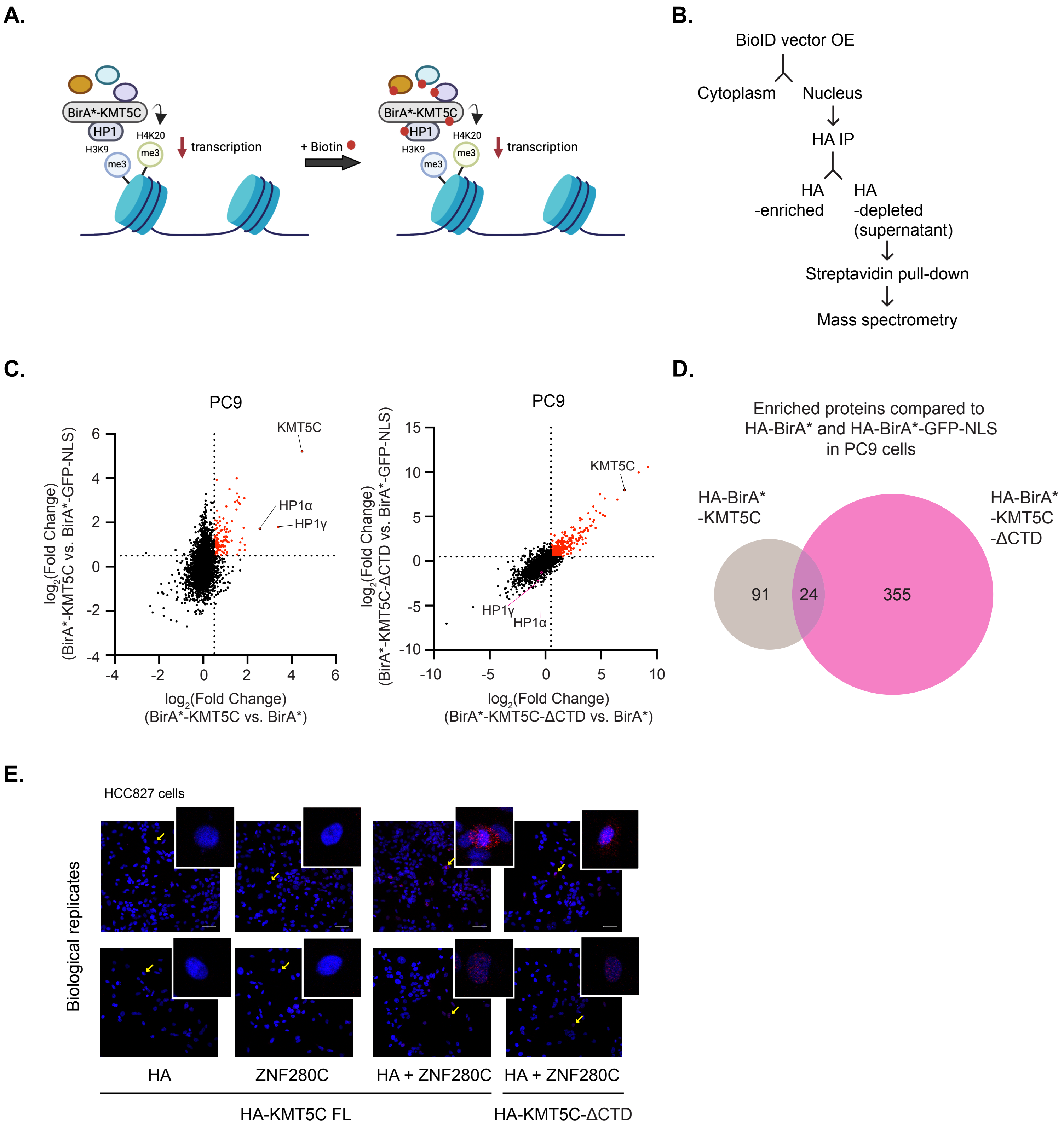
